# Raven: a de novo genome assembler for long reads

**DOI:** 10.1101/2020.08.07.242461

**Authors:** Robert Vaser, Mile Šikić

**Affiliations:** Laboratory for Bioinformatics and Computational Biology, University of Zagreb, Faculty of Electrical Engineering and Computing, Zagreb, Croatia; Laboratory of AI in Genomics, Genome Institute of Singapore, A^*^STAR, Singapore

## Abstract

We present new methods for the improvement of de novo genome assembly from erroneous long-reads incorporated into a straightforward tool called Raven (https://github.com/lbcb-sci/raven). Raven maintains similar performance for various genomes and has accuracy on par with other assemblers which support third-generation sequencing data. It is one of the fastest options while having the lowest memory consumption on the majority of benchmarked datasets.

---

Sequencing technologies have come a long way, from tiny fragments at their infancy to large chunks obtainable today. The relentless advances in both length and accuracy continue to alleviate the puzzle-like reconstruction problem of the sequenced genome, as more repetitive structures can be resolved naturally. Amidst the excess of available state-of-the-art options for de novo genome assembly^1–6^, we present a fast, memory frugal, reliable, and easy to use tool called Raven. It is an overlap-layout-consensus based assembler which accelerates the overlap step, builds an assembly graph^4^ from reads that were pre-processed with pile-o-grams^7^, implements a novel and robust simplification method based on graph drawings, and polishes the unambiguous graph paths with Racon^8^, all of which is compiled into a single executable.

Short substring matching is a conventional approach for similarity search in bioinformatics^9,10^. However, even with minimizers^4^ the overlap step of de novo assembly can take a substantial amount of time when handling larger genomes. To tackle this problem we enhanced the minimap^4^ algorithm following the MinHash approach^11^, where we select a fixed number of lexicographically smallest minimizers as the sequence sketch. The combination of MinHash on top of minimizers was already explored within the sequence mapper MashMap^12^, while a similar idea with hierarchical minimizers is the core of de novo assembler Peregrine^13^. Based on empirical evaluations, we opted for retaining |*read*|/*k* minimizers per read, where *k* is the minimizer length. Without any other algorithmic modifications to minimap, we are able to identify contained reads and create pile-o-grams for read pre-processing in a fraction of time and with a small impact on sensitivity. Suffix-prefix overlaps needed for graph constructions are found with the unmodified minimap algorithm within the containment-free read set, which is usually smaller than the whole sequencing yield by almost an order of magnitude.

Raven loads the whole sequencing sample into memory in compressed form, and finds overlaps in fixed-size blocks to decrease the memory footprint. Found overlaps are immediately transformed into pile-o-grams and discarded, except the longest few per read which are used for containment removal. Chimeric reads are iteratively identified and chopped by detecting sharp declines of coverage in pile-o-grams using coverage medians inferred from the stored overlaps. As minimap ignores the most frequent minimizers, which are critical for good repeat annotations, we lower this threshold while overlapping all contained reads to the set of containment-free reads, and search the updated pile-o-grams for sharp coverage inclines followed by sharp declines, both above the coverage median. Afterwards, the containment-free read set is overlapped to itself and repeat annotations are used to remove false overlaps between reads containing repetitive regions. Once the assembly graph is created, it is simplified stepwise with transitive reduction, tip removal, and bubble popping. Eventually, we simplify the graph with a novel method which lays out the graph in a two-dimensional Euclidean system, searches for edges that connect distant parts of the graph and removes them. Applying the force-directed placement algorithm^14^, which draws tightly connected vertices together, we can distinguish undetected chimeric or repeat-induced edges which are elongated with respect to others due to their rareness (Figure 1). Collapsing unambiguous paths while leaving room near junction vertices, coupled with the hierarchical force-calculation algorithm^15^, makes this drawing based simplification method feasible for even the largest assembly graphs. To finalize the assembly, contiguous paths of the graph are passed to two rounds of Racon.

**Figure 1.**
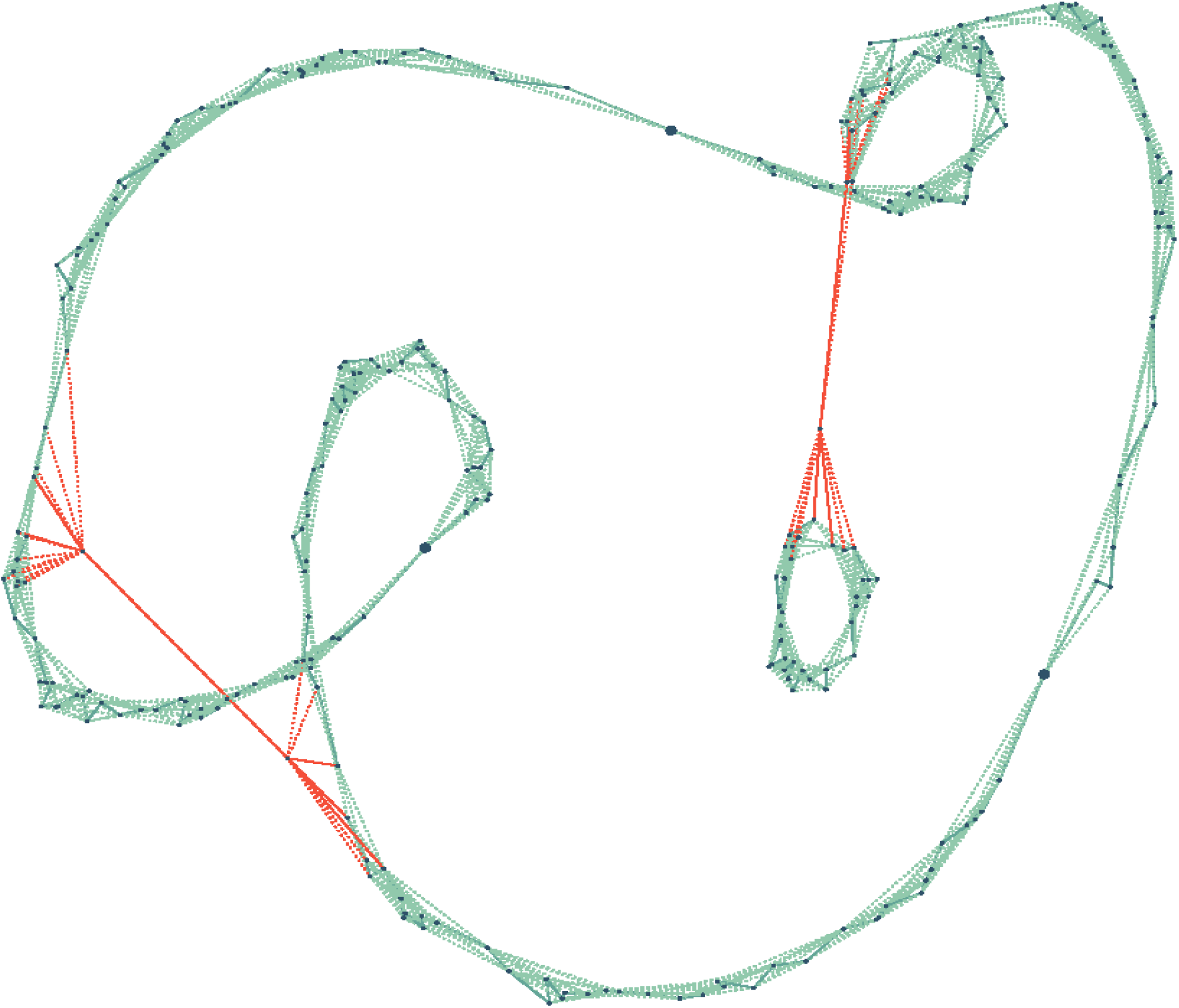
Bacterial assembly graph drawn with the force-directed placement algorithm. Raven uses vertex distances in two-dimensional Euclidean system to find elongated edges (red) that connect junction vertices and removes the longest ones. Those represent false connections which occur either due to sequencing errors or repetitive genomic regions. Without unitig creation (large circles) and the hierarchical force calculation, the drawing algorithm would partake an extensive amount of time on larger genomes. In addition, transitive edges (dotted green) are reinstated to increase the connectivity of neighboring vertices.

Since an earlier version of Raven proved as one of the best performers in a comprehensive benchmark^16^ at prokaryotic level, we evaluated several state-of-the-art assemblers alongside Raven on five model eukaryote datasets (Table 1), obtained by third-generation sequencing technologies, namely Pacific Biosciences (PacBio) and Oxford Nanopore Technologies (ONT). Emergence of PacBio’s High-Fidelity sequencing protocol (HiFi), and novel assemblers^13,17,18^ suitable for its highly accurate data, led us to evaluate the assembly reconstruction prospects of different sequencing approaches, that is ONT, Pacbio CLR (continuous long reads) and Pacbio HiFi, on three human samples (Table 2). Alongside default assembly quality metrics such as NGAx, genome fraction and accuracy, we evaluated gene completeness (single and multi-copy genes present both in the reference and the assembly), and where possible, the number of bacterial artificial chromosomes (BAC) resolved in an assembly. Details about computational cost can be found in the Supplementary (Table S1).

**Table 1.**
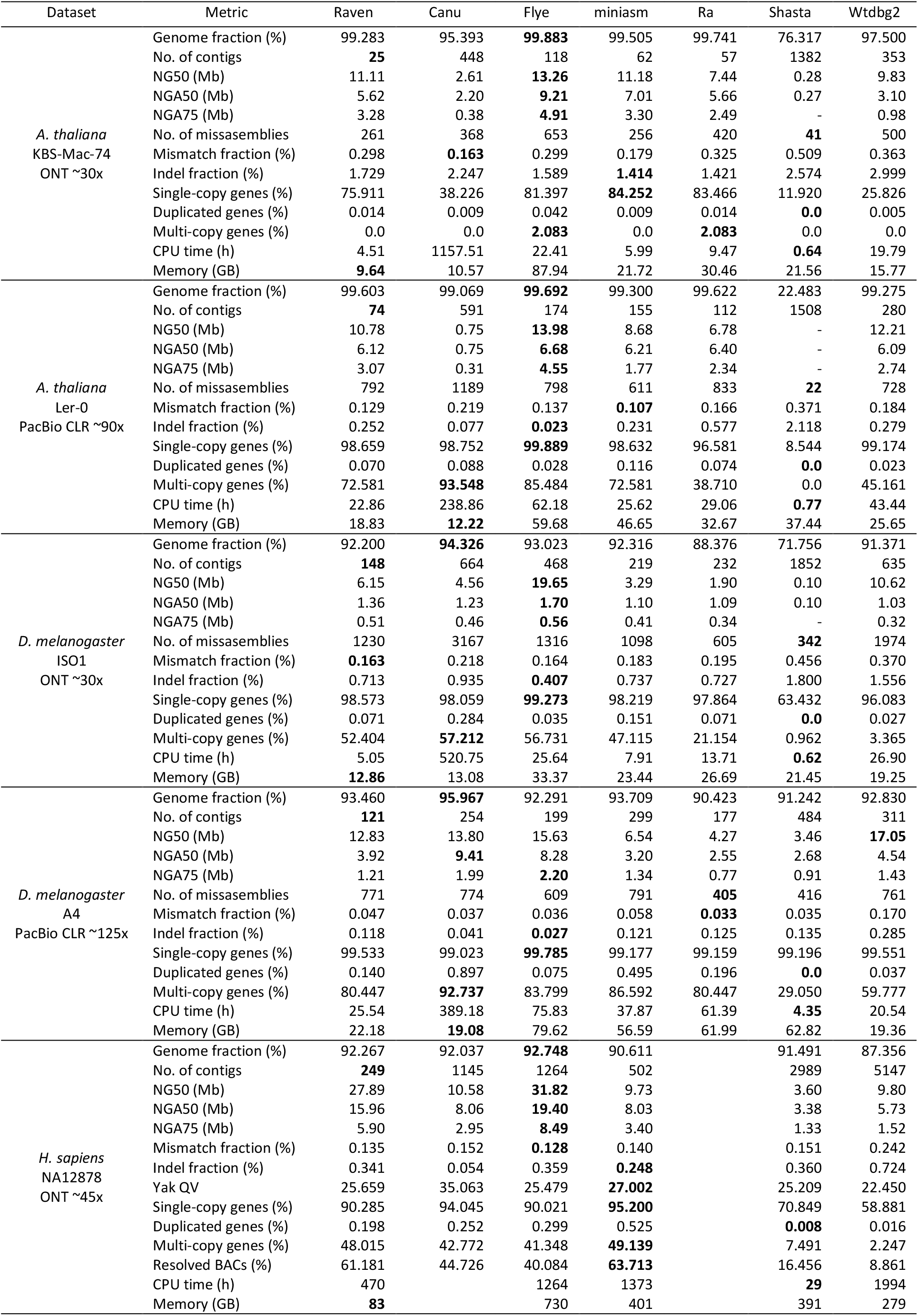
Evaluation of long-read assemblers.

**Table 2.**
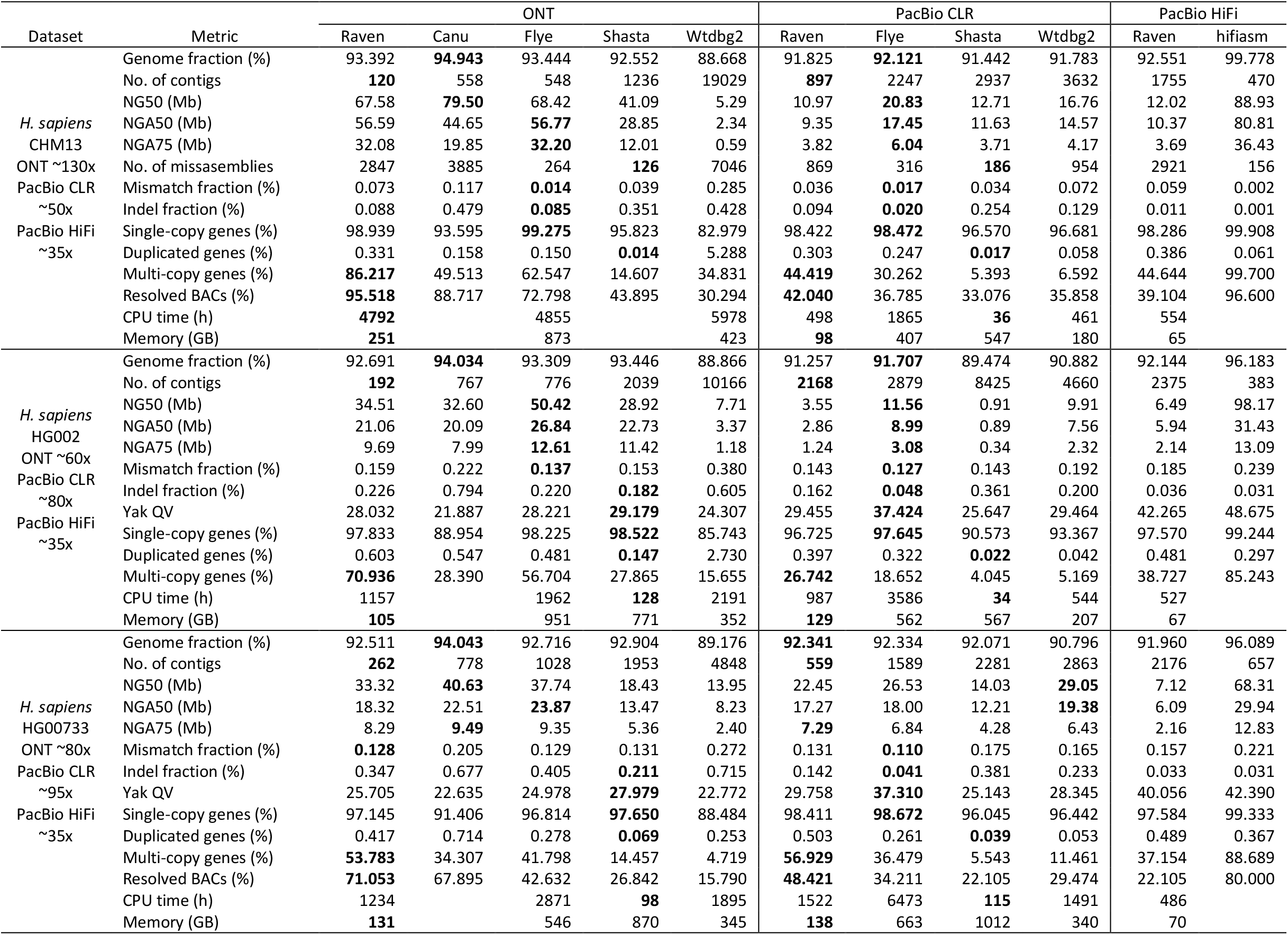
Evaluation of long-read assemblers across sequencing technologies.

On erroneous data, Raven is one of the fastest assemblers, and uses the least amount of memory on all but two datasets, while having better or comparable contiguity and accuracy. It especially stands out in the number of contigs with similar genome reconstruction fractions, and in the number of retained multy-copy genes and resolved BACs on human datasets. On the other hand, Raven does not utilize the accuracy of HiFi reads, which results in longer running times and subpar assembly results on more accurate data. We believe that more carefully tweaked parameters for the overlap step will lead to performance improvements.

We also run Raven on a couple of ONT plant datasets from two scientific studies^19,20^ and compared their results (Table 3). On datasets *B. oleracea, B. rapa* and *M. schizocarpa* Raven produces comparable assemblies to those obtained with Ra^21^. Furthermore, both *O. sativa* assemblies are more contiguous than the ones reported with Flye, but the BUSCO^22^ scores are lower as we did not polish our assemblies with Illumina data.

**Table 3.** Raven plant assemblies. Values in brackets represent assembly metrics in corresponding publications. Oryza genomes in the original publication were additionally polished with Illumina reads.

| Metric \ Dataset | *Brassica oleracea* | *Brassica rapa* | *Musa schizocarpa* | *Oryza sativa basmati 334* | *Oryza sativa dom sufid* |
| --- | --- | --- | --- | --- | --- |
| Total length (Mb) | 535.9 (546.4) | 351.7 (375.3) | 534.4 (522.0) | 382.4 (386.6) | 380.5 (383.6) |
| N50 (Mb) | 6.35 (7.28) | 5.52 (3.80) | 2.48 (2.13) | 8.14 (6.32) | 11.86 (10.53) |
| No. of contigs | 252 (244) | 410 (544) | 546 (615) | 116 (188) | 107 (116) |
| % complete BUSCOs | 74.783 (74.300) | 85.936 (79.700) | 47.150 (53.800) | 92.503 (97.600) | 92.193 (97.000) |
| CPU time (h) | 40.52 (261.40) | 58.55 (315.70) | 94.95 (245.60) | 43.59 (N/A) | 33.89 (N/A) |

Presented results indicate that PacBio HiFi assemblers achieve better overall reconstruction metrics, although ONT assemblies do not fall far off. ONT sequencing is still more approachable due to affordable consumables and portable devices, while requiring less gDNA than regular PacBio protocols. In addition, the length advantage of ONT reads and the recent increase in accuracy with the newest version of the Bonito basecaller (still in testing phase) justify the usage of assemblers which support this technology.

We showcased new algorithms for the overlap and layout phases of de novo genome assembly that reduce execution time and increase contiguity of the final assembly. We integrated them with an overlap module based on minimap, and the consensus module Racon, into a powerful standalone tool called Raven which is optimized for error-prone long reads. We argue that its performance coupled with the reduced cost per base of long-read sequencing technologies will enable assembly of large genomes even to laboratories with limited funding.

## Methods

Raven starts the assembly by constructing pile-o-grams (one-dimensional structures storing per-base coverage) and removing contained reads with the minimap algorithm, using 15-mers, a sliding window of 5 bases and discarding 10^−3^ most frequent minimizers. The whole sequencing data set is loaded into memory, replacing nucleotides with two bits and merging 64 succeeding Phred quality scores with their average. Reads are overlapped to each other in 1Gbp vs 4Gbp chunks, and only the lexicographically smallest |*read*|/15 minimizers are picked in both the index and the query (Supplementary Figure S1-S3; accuracy comparison in Supplementary Table S2). Once a block is processed, all overlaps are stacked into pile-o-grams which are decimated to every 16-th base. The longest 16 overlaps per read are stored for containment removal and connected component retrieval. When all pairwise overlaps are obtained, coverage medians are calculated for each pile-o-gram, reads are trimmed to the longest region covered with at least 4 other reads, and potential chimeric sites are detected by finding bases which have 1.82 times smaller coverage than their neighboring bases. Contained reads are dropped only if the containing read does not have a potential chimeric region. Decreasing the number of reads through containment removal enables faster verification of chimeric annotations. Given the stored suffix-prefix overlaps, Raven finds connected components and their coverage median, which approximates the sequencing depth. Each annotated coverage drop is used to chop problematic reads to their longest non-chimeric region, if the drop is consistent with the coverage median of the connected component the read belongs to. The whole process is done iteratively to capture different molecule copy numbers, because resolving chimeric reads tends to the forming of new connected components. Another containment check is carried out once chimeric sequences are resolved.

Afterwards, Raven searches for suffix-prefix overlaps between the remaining reads enforcing the use of all minimizers. In addition, all contained reads are overlapped to the containment-free read set in order to increase the coverage of repetitive regions, again employing the MinHash approach. Decreasing the minimizer frequency filter to 10^−5^ enables proper repeat annotation in which sought bases need to have coverage at least 1.42 times larger than the component coverage median. Repetitive regions at either end of a read are used to iteratively remove false overlaps, i.e. overlaps that connect different copies of bridged repeats (repetitive genomic regions that are entirely contained in at least one read).

Once the overlap set is cleaned, the assembly graph is built and simplified stepwise with standard layout algorithms such as transitive reduction, tipping, and bubble popping. Information about transitive connections is kept for the last simplification step, which plots the assembly graph in a two-dimensional space, in order to increase the connections between neighboring vertices. Raven searches for edges connecting remote parts of the graph, which are usually present due to leftover sequencing artefacts or unresolved repeats. The force-directed placement algorithm enlarges most of such edges due to their rareness. Given the quadratic time complexity *0(*|*V*|^2^)^14^ and an approximate of 100 iterations until convergence, we shrink the graph by creating unitigs (paths in the graph consisting of vertices with only one ingoing and one outgoing edge) that are 42 vertices away from any junction vertex (vertices with more than one outgoing or ingoing edge). Furthermore, approximating the forces of distant vertices by replacing them with their centre of mass enables linearithmic time complexity *0(*|*V*| log|*V*|)^15^, and the use of this method on larger genomes. Depending on vertex distances in a finished drawing, Raven removes outgoing edges that are at least twice as long as any other outgoing edge of that junction vertex. As the drawing heavily depends on an initial layout, which is random but with a fixed seed, the whole procedure is restarted 16 times. It should be noted that if there exist a lot of false connections in a single area of the graph (usually induced by repeats), the drawing algorithm will not be able to sufficiently enlarge all of these edges for removal (Supplementary Figure S4).

Finally, paths of the assembly graph without external branches are polished with a library version of Racon, using small windows of 500bp and partial order alignment with linear gaps, in a total of two iterations. All constant values used in various Raven stages were empirically determined based on a large set of real datasets of various sizes.

Because of resource limitations we chose the best performing genome assemblers for erroneous third-generation data from recent scientific papers^3,6,16^. The assemblers are Raven (v1.3.0), Canu (v2.0), Flye (v2.8.1), miniasm (v0.3-r179) coupled with minimap (v0.2-r123) and polished with two iterations of Racon (v1.4.13), Ra (v0.2.1), Shasta (v0.7.0) and Wtdbg2 (v2.5). Raven was run without any additional parameters on ONT and PacBio CLR datasets. On PacBio HiFi datasets, we increase k-mer length from 15 to 29, and window length from 5 to 9, in order to decrease the number of found pairwise overlaps (comparison with default parameters can be found in Supplementary Table S3). We use options ‘-pacbio’ or ‘-nanopore’ for Canu, ‘-pacbio-raw’ or ‘-nano-raw’ for Flye, ‘-x ont’ or ‘-x pb’ for Ra, ‘-x sq’, ‘-x rs’ or ‘-x ont’ for Wtdbg2, and configuration files Nanopore-Dec2019, Nanopore-Sep2020 or PacBio-CLR-Dec2019 for Shasta. For ONT runs we modified the Shasta consensus caller to better match the basecaller used to obtain the corresponding dataset, while we decreased the minimal read length to 5000 for non-human datasets, except PacBio CLR *D. melanogaster* dataset for which Shasta produced a decent assembly. Canu and Wtdbg2 require approximate genomes sizes which were 120 Mb, 144 Mb, and 3 Gb for *A. thaliana, D. melanogaster* and *H. sapiens* datasets, respectively. All assemblers were run with 64 threads on a server with 1 TB RAM and two AMD EPYC™ 7702 64-core processors. Due to high memory requirements, the ONT CHM13 dataset was benchmarked with 48 threads on a server with two Intel^®^ Xeon^®^ Platinum 8260L 24-core processors and 1.5 TB of Optane™ Persistent Memory. Shasta was unable to assemble the PacBio CLR HG00733 dataset on the first machine due to memory requirements, so it was run on the second machine. Also, it was not able to assemble the ONT CHM13 dataset on either machine, so we found the assembly in its publication. Canu was not run on human datasets due to its long running time, but we found assemblies in other publications^5,23^ (NA12878 assembly was polished with Illumina data so it was excluded from accuracy comparison). We omitted Ra from the human dataset benchmark due to its complexity on larger genomes. Hifiasm human assemblies were found in its publication^18^.

We used QUAST-LG^24^ (v5.0.2) for assembly evaluation and ran it with minimal identity of 80%. For *H. sapiens* datasets we used the T2T (telomere-to-telomere) reconstruction of CHM13 (and options ‘--large’ in QUAST), while for *A. thaliana* and *D. melanogaster* datasets we used appropriate NCBI assemblies or references depending on the strain. The assembly quality value (QV) was obtained with yak (v0.1), which is available at https://github.com/lh3/yak, by comparing 31-mers found in short accurate reads and the assembly for datasets NA12878, HG002 and HG00733. Gene completeness was evaluated with paftools (v2.17-r982) asmgene function, found inside the minimap2^25^ package. We mapped annotated Ensembl cDNA sequences (v102 for *D. melanogaster* and *H. sapiens*, and v49 for *A. thaliana*) to the references and the assemblies. Identity of 97% was used to find single-copy and duplicated single-copy genes, while 99% identity was used for multi-copy genes. We validated BAC resolution with a pipeline available at https://github.com/skoren/bacValidation (commit 4f3e463), where 99.5% of bases of a BAC need to be present in the assembly for it to be resolved. We used VMRC53 (237 BACs), VMRC59 (647 BACs) and VMRC62 (190 BACs) clones for NA12878, CHM13 and HG00733, respectively. BUSCO (v4.1.4) scores for the five plant datasets were found with the *embryophyta* database, although the current version contains more orthologs (1614 in total).

ONT dataset for *A. thaliana* is available under the accession number ERR2173373, for *D. melanogaster* under SRR6702603, for *H. sapiens* NA12878 [here](https://github.com/nanopore-wgs-consortium/NA12878) (release 6), for *H. sapiens* CHM13 [here](https://github.com/nanopore-wgs-consortium/CHM13) (release 6), for *H. sapiens* HG002 [here](https://ftp-trace.ncbi.nlm.nih.gov/giab/ftp/data/AshkenazimTrio/HG002_NA24385_son/Ultralong_OxfordNanopore/guppy-V3.4.5/), and for *H. sapiens* HG00733 [here](https://s3-us-west-2.amazonaws.com/human-pangenomics/index.html?prefix=NHGRI_UCSC_panel/HG00733/nanopore/).

PacBio CLR dataset for *A. thaliana* is available [here](https://downloads.pacbcloud.com/public/SequelData/ArabidopsisDemoData/), for *D. melanogaster* under accession number SRR5439404, for *H. sapiens* CHM13 [here](https://github.com/nanopore-wgs-consortium/CHM13) (extracted from draft v1.0 bam), for *H. sapiens* HG002 [here](https://ftp-trace.ncbi.nlm.nih.gov/giab/ftp/data/AshkenazimTrio/HG002_NA24385_son/PacBio_MtSinai_NIST/PacBio_fasta/), and for *H. sapiens* HG0073 under SRR7615963.

PacBio HiFi dataset for *H. sapiens* CHM13 is available from accession number SRR11292120 to SRR11292123, for *H. sapiens* HG002 under SRR10382244, SRR10382245, SRR10382248 and SRR10382249, and for *H. sapiens* HG00733 under ERX3831682.

Illumina reads for yak evaluation are available from accession number SRX1049768 to SRX1049782 for *H. sapiens* NA12878, [here](https://ftp-trace.ncbi.nlm.nih.gov/giab/ftp/data/AshkenazimTrio/HG002_NA24385_son/NIST_HiSeq_HG002_Homogeneity-10953946/NHGRI_Illumina300X_AJtrio_novoalign_bams/) (extracted from 60x bam) for *H. sapiens* HG002, and under accession number SRR7782677 for *H. sapiens* HG00733.

Accession numbers of the plant datasets used for separate Raven evaluation can be found in corresponding publications.

All generated assemblies in this research can be found at Zenodo under DOI 10.5281/zenodo.4443062.

## Acknowledgments

This work has been supported in part by the Croatian Science Foundation under the project Single genome and metagenome assembly (IP-2018-01-5886), by the European Regional Development Fund under the grant KK.01.1.1.01.0009 (DATACROSS), and by the A*STAR Computational Resource Centre through the use of their high-performance computing facilities. R.V. and M.Š. have been partially supported by funding from A*STAR, Singapore. We acknowledge Intel® Corporation for allowing us to test with Intel® Optane™ Persistent Memory server and providing us with high-quality technical support. Finally, we thank Goran Žužić from Carnegie Mellon University for useful discussions in the field of graph drawings.

## Notes

https://github.com/lbcb-sci/raven

### Competing Interest Statement

The authors have declared no competing interest.

### Summary of Updates

We have updated the assembly benchmark in the manuscript with additional human datasets covering different sequencing techniques, included newer versions of assemblers, and conducted additional assembly evaluation methods. We polished the manuscript, added missing references, emphasized best values in tables, and wrote a supplementary.

