## Supplementary for "Raven: a de novo genome assembler for long reads"

### Supplementary information

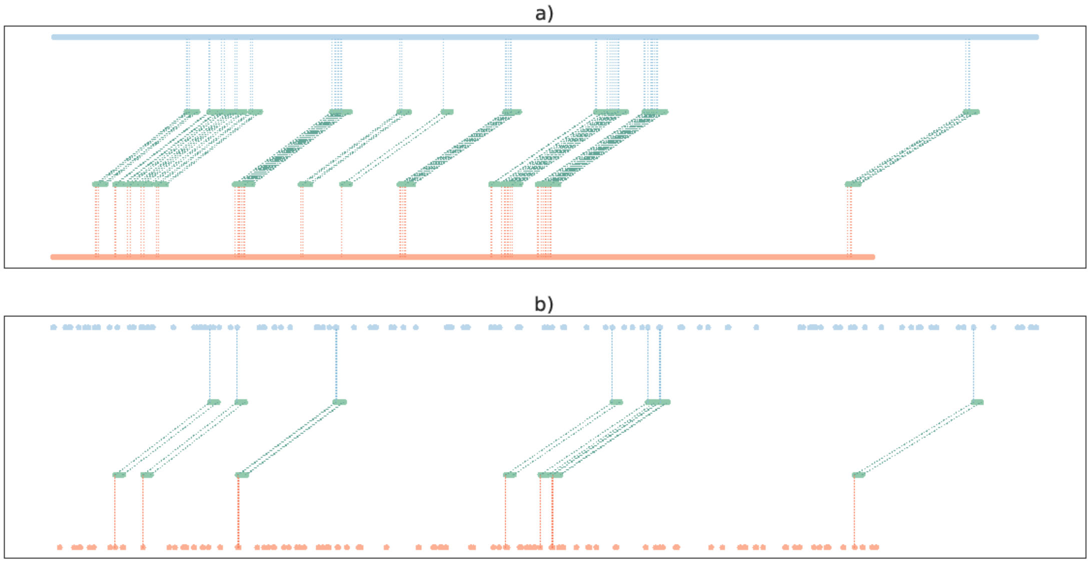


**Figure S1 Overlap between two erroneous reads based on minimizer matches.** Raven uses the minimap algorithm to find pairwise overlaps, in which lexicography smallest k-mers in sliding windows (minimizers) of both reads are collected (blue and orange) and a linear chain of matches is found (green). a) While collecting all minimizers from a small sliding window ensures the retrieval of most overlaps between similar reads, b) a decent amount of overlaps can be retained by picking only a portion of the smallest minimizers. Shrinking the minimizer search space, without any other modifications, greatly accelerates the algorithm, and justifies the impact on sensitivity for containment removal and pile-o-gram creation.

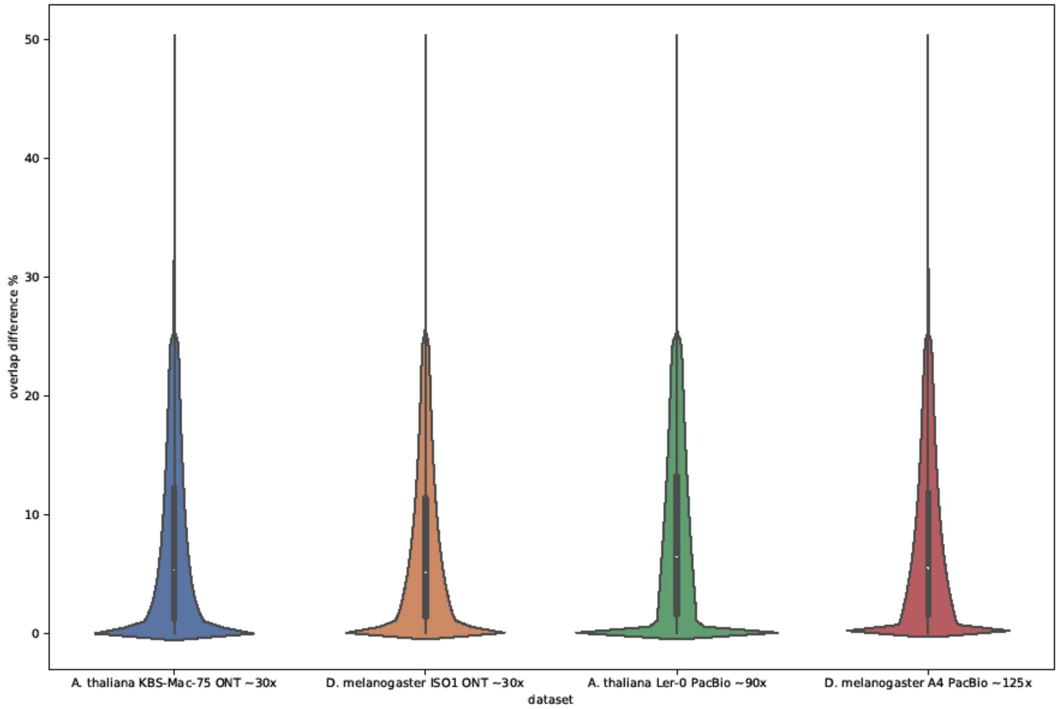


**Figure S2 Overlap difference distributions when employing the MinHash paradigm on top of minimizers.** Raven reduces the amount of minimizers used in the minimap algorithm by choosing only a portion of smallest values per read, which affects the beginning and ending position of pairwise overlaps between reads, but enables faster containment removal with a small sensitivity degradation. Depicted values represent the absolute difference between old and new coordinates divided by the old overlap length (we ignore overlaps which cover less than 75% of original region on either of the reads).

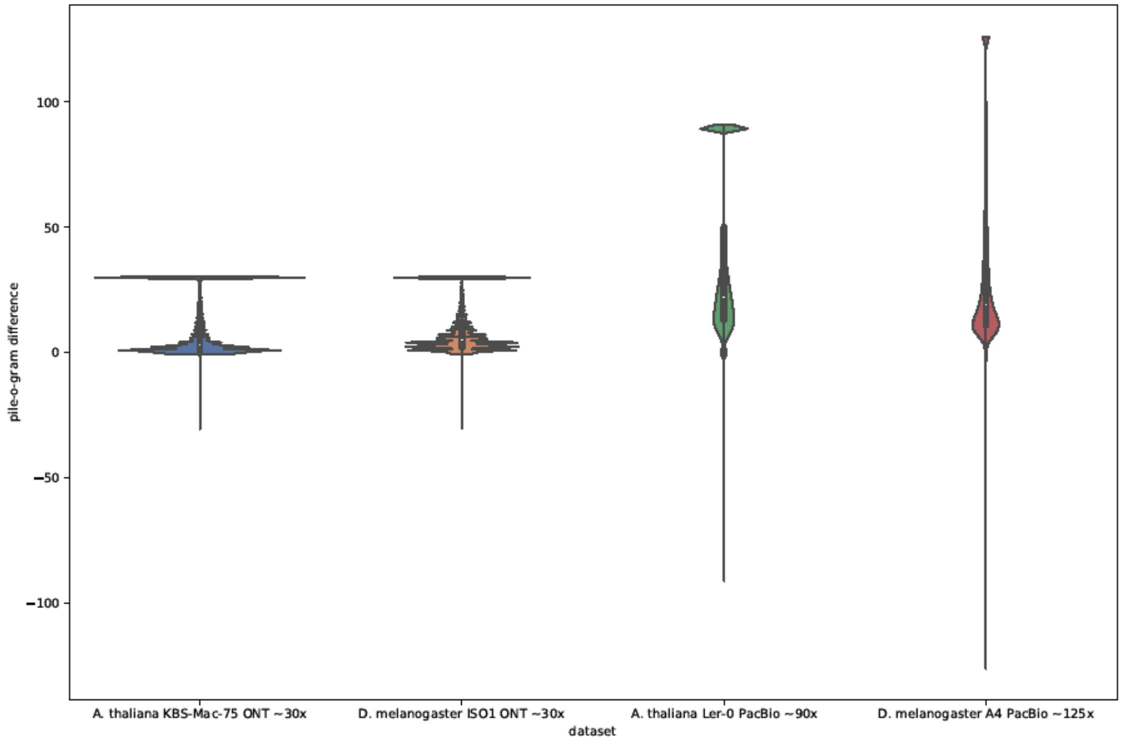


**Figure S3 Pile-o-gram difference distribution when employing the MinHash paradigm on top of minimizers.** Raven reduces the amount of minimizers used in the minimap algorithm by choosing only a portion of smallest values per read, which affects per-base coverage in pile-o-grams, but it is negligible for chimeric and repeat annotations. Depicted values represent per-base differences between old and new pile-o-grams, which are saturated to the sequencing depth in absolute value (the biggest differences in coverage are in repetitive regions due to different minimizer sampling).

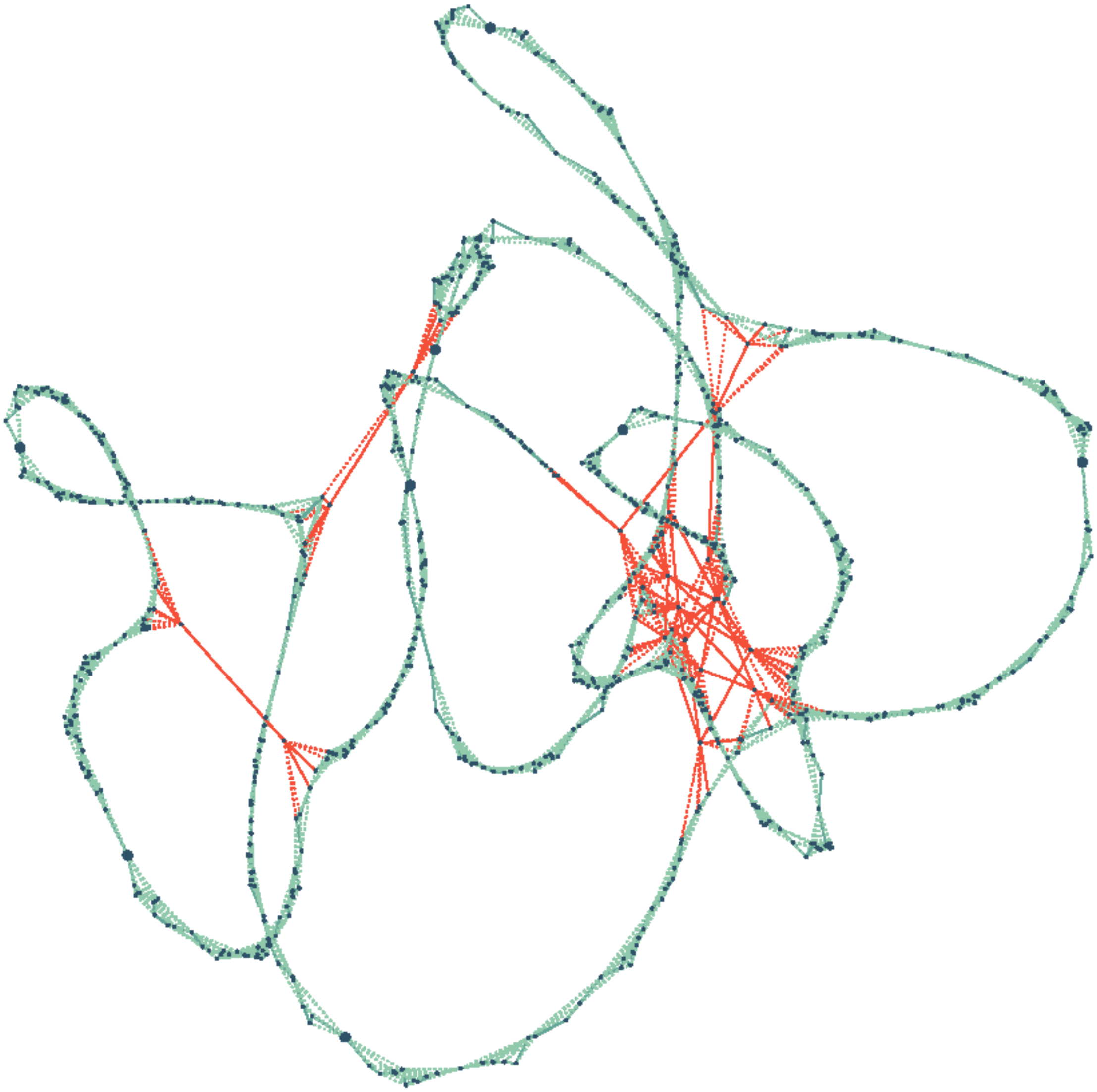

**Figure S4 Bacterial assembly graph constructed without read pre-processing and drawn with the force-directed placement algorithm.** The graph simplification method based on vertex distances in two-dimensional Euclidean system underperforms when there are many repeat induced edges (red) which connect distant genomic regions. Compared to Figure 1, we skipped overlap removal based on repeat annotations, which yields a more tangled graph at the end of the layout phase. Although, the simplification method is still able to correctly resolve all false connections for this case, but in a total of two iterations. Number of edges removed this way constitutes a small portion of edges in the assembly graph and depends on many intertwined factors (for *A. thaliana* and *D. melanogaster* datasets this value ranges between 1% and 11% with respect to number of remaining edges at the end of the layout phase). To fully utilize this method, read pre-processing should be applied beforehand.

**Table S1 Estimates of assembly cost.** Values with asterisks were found in publications of corresponding assemblers, while values with tilde are approximations when more threads are invoked. Canu assemblies were omitted due to long running times (according to (Shafin et al. 2020), its estimate cost is around $19000 for all three ONT datasets).

| Dataset | Assembler | Number of threads | Real Time (h) | Memory (Gb) | AWS instance | AWS cost / hour ($) | AWS cost ($) |
|---|---|---|---|---|---|---|---|
| CHM13 ONT ~130x | Raven | 48 | 123.12 | 251 | m5a.16xlarge | 2.75 | ~253.94 |
| | Flye | 48 | 141.78 | 873 | x1.16xlarge | 6.67 | ~709.25 |
| | Shasta | 128* | 5.28* | | x1.32xlarge | 13.34 | 70.43 |
| | Wtdbg2 | 48 | 128.58 | 423 | r5a.16xlarge | 3.62 | ~349.09 |
| HG002 ONT ~60x | Raven | 64 | 25.21 | 105 | c5a.16xlarge | 2.46 | 62.02 |
| | Flye | 64 | 49.86 | 951 | x1.16xlarge | 6.67 | 332.57 |
| | Shasta | 64 | 3.09 | 771 | x1.16xlarge | 6.67 | 20.61 |
| | Wtdbg2 | 64 | 36.59 | 352 | r5a.16xlarge | 3.62 | 132.46 |
| HG00733 ONT ~80x | Raven | 64 | 26.88 | 131 | m5a.16xlarge | 2.75 | 73.92 |
| | Flye | 64 | 57.14 | 546 | x1.16xlarge | 6.67 | 381.12 |
| | Shasta | 64 | 2.83 | 870 | x1.16xlarge | 6.67 | 18.88 |
| | Wtdbg2 | 64 | 31.46 | 345 | r5a.16xlarge | 3.62 | 113.89 |
| CHM13 PB ~50x | Raven | 64 | 10.41 | 98 | c5a.16xlarge | 2.46 | 25.61 |
| | Flye | 64 | 35.29 | 407 | r5a.16xlarge | 3.62 | 127.75 |
| | Shasta | 64 | 1.54 | 547 | x1.16xlarge | 6.67 | 10.27 |
| | Wtdbg2 | 64 | 8.06 | 180 | m5a.16xlarge | 2.75 | 24.64 |
| HG002 PB ~80x | Raven | 64 | 23.78 | 129 | m5a.16xlarge | 2.75 | 65.40 |
| | Flye | 64 | 63.50 | 562 | x1.16xlarge | 6.67 | 423.55 |
| | Shasta | 64 | 1.50 | 567 | x1.16xlarge | 6.67 | 10.01 |
| | Wtdbg2 | 64 | 9.29 | 207 | m5a.16xlarge | 2.75 | 25.55 |
| HG00733 PB ~95x | Raven | 64 | 30.92 | 138 | m5a.16xlarge | 2.75 | 85.03 |
| | Flye | 64 | 110.94 | 663 | x1.16xlarge | 6.67 | 800.00 |
| | Shasta | 48 | 1.28 | 1012 | x1.32xlarge | 13.34 | ~6.40 |
| | Wtdbg2 | 64 | 25.12 | 340 | r5a.16xlarge | 3.62 | 90.93 |
| HiFi ~35x | hifiasm | 48* | 9* | 150* | m5a.12xlarge | 2.06 | 18.54 |

**Table S2 Sensitivity and execution time impact of minimizer and MinHash combination.**

| | *A. thaliana* KBS-Mac-75 ONT ~30x | *D. melanogaster* ISO1 ONT ~30x | *A. thaliana* Ler-0 PacBio ~90x | *D. melanogaster* A4 PacBio ~125x |
|---|---|---|---|---|
| No. of overlaps - minimizers | 33639084 | 38522454 | 250456231 | 376494493 |
| No. of overlaps - minimizers and MinHash | 8724876 | 15282409 | 26442472 | 153956450 |
| Overlap Jaccard score | 0.208 | 0.238 | 0.057 | 0.266 |
| Containment Jaccard score | 0.800 | 0.835 | 0.698 | 0.864 |
| Speedup | 4.42 | 3.62 | 4.21 | 4.01 |

**Table S3 Impact of overlap parameters (k-mer length and sampling window length) on Raven's performance on HiFi data.**

| | *H. sapiens* CHM13 HiFi ~35x | | *H. sapiens* HG002 HiFi ~35x | | *H. sapiens* HG00733 HiFi ~35x | |
|---|---|---|---|---|---|---|
| Metric | Raven (15,5) | Raven (29,9) | Raven (15,5) | Raven (29,9) | Raven (15,5) | Raven (29,9) |
| Genome fraction (%) | 91.492 | 92.551 | 91.199 | 92.144 | 91.135 | 91.960 |
| No. of contigs | 5689 | 1755 | 7282 | 2375 | 7615 | 2176 |
| NG50 (Mb) | 1.05 | 12.02 | 0.83 | 6.49 | 0.77 | 7.12 |
| NGA50 (Mb) | 0.80 | 10.37 | 0.63 | 5.94 | 0.57 | 6.09 |
| NGA75 (Mb) | 0.33 | 3.69 | 0.25 | 2.14 | 0.23 | 2.16 |
| No. of missasemblies | 3803 | 2921 | 5513 | 4554 | 5389 | 3743 |
| Mismatch fraction (%) | 0.042 | 0.059 | 154.54 | 0.185 | 0.142 | 0.157 |
| Indel fraction (%) | 0.009 | 0.011 | 34.17 | 0.036 | 0.034 | 0.033 |
| Single-copy genes (%) | 95.014 | 98.286 | 93.967 | 97.570 | 94.173 | 97.584 |
| Duplicated genes (%) | 1.905 | 0.386 | 1.817 | 0.481 | 1.600 | 0.489 |
| Multi-copy genes (%) | 42.097 | 44.644 | 36.929 | 38.727 | 33.258 | 37.154 |
| Resolved BACs (%) | 28.594 | 39.104 | | | 18.421 | 22.105 |

| | | | | | | |
|---|---|---|---|---|---|---|
| CPU time (h) | 1313 | 554 | 3449 | 527 | 1300 | 486 |
| Memory (GB) | 87 | 65 | 91 | 67 | 97 | 70 |
